# The Iron Content of Human Serum Albumin Modulates the Susceptibility of *Acinetobacter baumannii* to Cefiderocol

**DOI:** 10.1101/2022.08.24.505215

**Authors:** Jenny Escalante, Brent Nishimura, Marisel R. Tuttobene, Tomás Subils, Vyanka Mezcord, Luis A. Actis, Marcelo E. Tolmasky, Robert A. Bonomo, María Soledad Ramirez

## Abstract

Mortality rates of patients infected with *Acinetobacter baumannii* treated with cefiderocol (CFDC) were not as favorable as the best available treatment for pulmonary and bloodstream infections. Previous studies showed that the presence of human serum albumin (HSA) or HSA-containing fluids like human pleural fluid (HPF) or human serum (HS) in the growth medium is correlated with a decrease in the expression of genes associated with high-efficiency iron uptake systems. These observations may explain the less-than-ideal performance of CFDC in pulmonary and bloodstream infections because ferric siderophore transporters enhance penetration of CFDC into the cell’s cytosol. Removal of HSA from HPF or HS resulted in a reduction of the minimal inhibitory concentration of CFDC. Concomitant with these results, there was an enhancement of the expression of genes associated with high-efficiency iron uptake systems. In addition to inducing modifications in iron-uptake gene expression, removal of HSA also decreased the expression of β-lactam resistance genes. Taken together, these observations indicate that environmental HSA has a role in the expression levels of selected *A. baumannii*. Furthermore, removal of iron from HSA had the same effect as removal of HSA on the expression of genes associated with high-efficiency iron uptake systems, suggesting that at least one of the mechanisms by which HSA regulates the expression of selected genes is through acting as an iron supplier.

**IMPORTANCE:** Cefiderocol (CFDC) is a new antibiotic that combines its major bactericidal activity, i.e., inhibition of the Gram-negative bacterial cell wall synthesis, with a first in its class mechanism of cell penetration. The siderophore-like moiety facilitates entry through receptors that recognize ferric-siderophore complexes. Recent trials showed that treating pulmonary and bloodstream *Acinetobacter baumannii* infections with CFDC did not result in the same outcomes as treating other pathogens. Our studies indicated that exposure to human fluids that contain human serum albumin (HSA) increases the MIC values of CFDC. Results described in this work show that HSA is responsible for a reduction in susceptibility of *A. baumannii* to CFDC. Furthermore, the presence of HSA in the milieu produces a reduction in levels of expression of proteins associated with high-affinity iron uptake systems and enhanced expression of β-lactam resistance-associated genes. Deferration of HSA was accompanied by a loss of the ability to modify these genes’ expression levels. These results indicate that the microbiological activity of CFDC towards *A. baumannii* is attenuated in the presence of HSA-containing fluids. This unique insight opens up new avenues of investigation. Understanding this phenomenon’s molecular mechanism will help define methodologies to increase treatment efficiency.

## INTRODUCTION

The development of novel and effective antibiotic treatments is an urgent need created by the increased number of antibiotic resistant bacteria. Multi- or pan-resistant *Acinetobacter baumannii* strains, recognized as urgent threats by the Centers for Disease Control and Prevention (CDC), are responsible for serious and often untreatable hospital-acquired infections (1, 2). Especially worrisome are infections caused by carbapenem-resistant *A. baumannii* (CRAB) strains for which only a few treatment options are still viable (3-5).

Cefiderocol (CFDC), a recently approved broad-spectrum antibiotic, is one of the few available options to treat CRAB infections (6-10). CFDC is a hybrid molecule that consists of a cephalosporin component, which targets cell wall synthesis, linked to a catechol moiety, which facilitates cell penetration by active ferric siderophore transporters (6-9, 11). CFDC has been approved to treat nosocomial pneumonia and urinary tract infections caused by extensively drug-resistant (XDR) Gram-negative bacteria (4, 12, 13). However, the CREDIBLE-CR randomized trial showed that the mortality rates of patients infected with *A. baumannii* and treated with CFDC were higher than the best available treatment for pulmonary and bloodstream infections (4). In contrast, mortality rates did not increase in CFDC-treated urinary tract infections (4).

Previous work showed that HSA and HSA-containing fluids, such as human pleural fluid (HPF) and human serum (HS), modulate the transcriptional expression of various genes associated with several *A. baumannii* functions, including antibiotic resistance, DNA acquisition, and iron uptake (14-17). The changes in the transcriptional expression of genes associated with iron uptake systems can partly explain the lower success seen in treating *A. baumannii* infection with CFDC (4, 18, 19).

The addition of HS, HPF, or purified HSA to the growth medium was associated with an increase in the CFDC MICs of three CRAB clinical isolates with different genetic backgrounds (20). These conditions also produced modifications in gene expression. Genes that are part of high-affinity iron-uptake systems were downregulated and those associated with resistance to β-lactams, such as *pbp1, pbp3, bla*_OXA-51-like_, *bla*_PER-7_, *bla*_ADC_, were upregulated (20). In contrast, the addition of human urine (HU), which contains only traces of HSA or free iron, did not result in modifications of CFDC MICs. Also, these conditions produced an enhancement of transcription of *piuA, pirA, bauA*, and *bfnH* (21). These results strongly suggest that human bodily fluids with high HSA content induce changes in the expression of iron uptake and β-lactam resistance-associated genes in *A. baumannii*. Modifications in the levels of expression of high-affinity iron uptake system components results in variations of CFDC MIC values. We hypothesize that the presence of HSA creates an iron-rich environment that represses the expression of iron-uptake genes limiting CFDC’s entrance into bacterial cells. In this work we analyze the effect of HSA on CFDC and on the expression of genes coding for siderophore-mediated iron acquisition functions in *A. baumannii*. We also show that at least one of the mechanisms by which HSA regulates transcription of genes coding for siderophore-mediated iron acquisition is associated with HSA acting as an iron carrier consequentially facilitating iron to *A. baumannii*.

## RESULTS AND DISCUSSION

### *HSA present in human fluids alters CFDC MICs and* A. baumannii *via a global transcriptional response*

HSA, the predominant protein in human plasma and extracellular fluids, acts as a key host signal triggering an adaptive response in a variety of pathogens and serves as an important host iron reservoir (14, 16, 20-26). Therefore, we sought to determine whether HSA causes changes in the MIC values of CFDC when *A. baumannii* is exposed to HPF or HS, host fluids that have 50-70 % and 90-96 % HSA, respectively. For this purpose, the *A. baumannii* CRAB strains AB5075 and AMA40 harboring *bla*_OXA-23_ and *bla*_NDM-1,_ respectively, were cultured in LB or LB in the presence of HPF or HS as well as in the presence of the cognate HSA-free derivatives of these two human fluids that were prepared as described in Material s and Methods.

This analysis showed a slight but decrease in the CFDC MIC for strain AB5075 when HSA was removed from both fluids (1 doubling dilution) (Table 1). For strain AMA40, the effect observed when HSA was not present was more pronounced. A decrease of 7 and 4 doubling dilutions was seen when HSA was removed from HPF and HS, respectively (Table 1). Notably, in addition, the appearance of colonies within the growth inhibition zones (intracolonies) was detected in samples cultured in the presence of HPF and HS but not when bacteria were cultured under HSA depleted conditions.

**Table 1:** Minimal Inhibitory Concentrations (MICs) of cefiderocol (CFDC) for the CRAB AB5075 and AMA40 strains, performed using CFDC MTS strips (Liofilchem S.r.l., Italy) on Iron-depleted CAMHA (Cation Adjusted Mueller Hinton Agar) and the different conditions tested.

| Condition | CFDC MIC (mg/L) |  |
| --- | --- | --- |
|  | AB5075 | AMA40 |
| CAMHB | 0.5 (S) | 0.5 (S) |
| 4% HPF | 1 (S) | 16* (R) |
| 4% HPF HSA free ** | 0.5 (S) | 0.25 (S) |
| 100% HS | 1 (S) | 4* (S) |
| 100% HS HSA free ** | 0.5 (S) | 0.25 (S) |

CFDC: cefiderocol, S: Susceptible, R: Resistant
\* Intra-colonies are present.
\*\*HSA Removal, Sigma Aldrich

To further confirm the role of HSA as a specific inducer of these *A. baumannii* responses, the transcriptomic analysis by quantitative RT-PCR (qRT-PCR) of both strains cultured in the presence of HS and HPF and their cognate HSA-free derivatives was also assessed. The expression of *bauA, pirA, piuA, bfnH, tonb3*, and *basE*, which code for active iron uptake functions, and genes coding for β-lactam resistance (*bla*_OXA-69_, *pbp3*, and *pbp1*), all of which are affected by the presence of HPF and HS (27), were evaluated. In both CRAB strains, we observed that the expression of the iron uptake genes was significantly increased when HSA was not present in HS (Figures 1A and Figure 2A, Table S1), suggesting that HSA plays a specific role triggering the observed changes. As predicted considering previous observation (27), an opposite result was observed when we analyzed the expression of the β-lactam resistance genes. A statistically significant decrease in the level of expression of *bla*_OXA-69_, *pbp3*, and *pbp1* was seen when these *A. baumannii* strains were incubated in the presence of HS lacking HSA (Figures 2A and 2B, Table S1).

**Figure 1.**
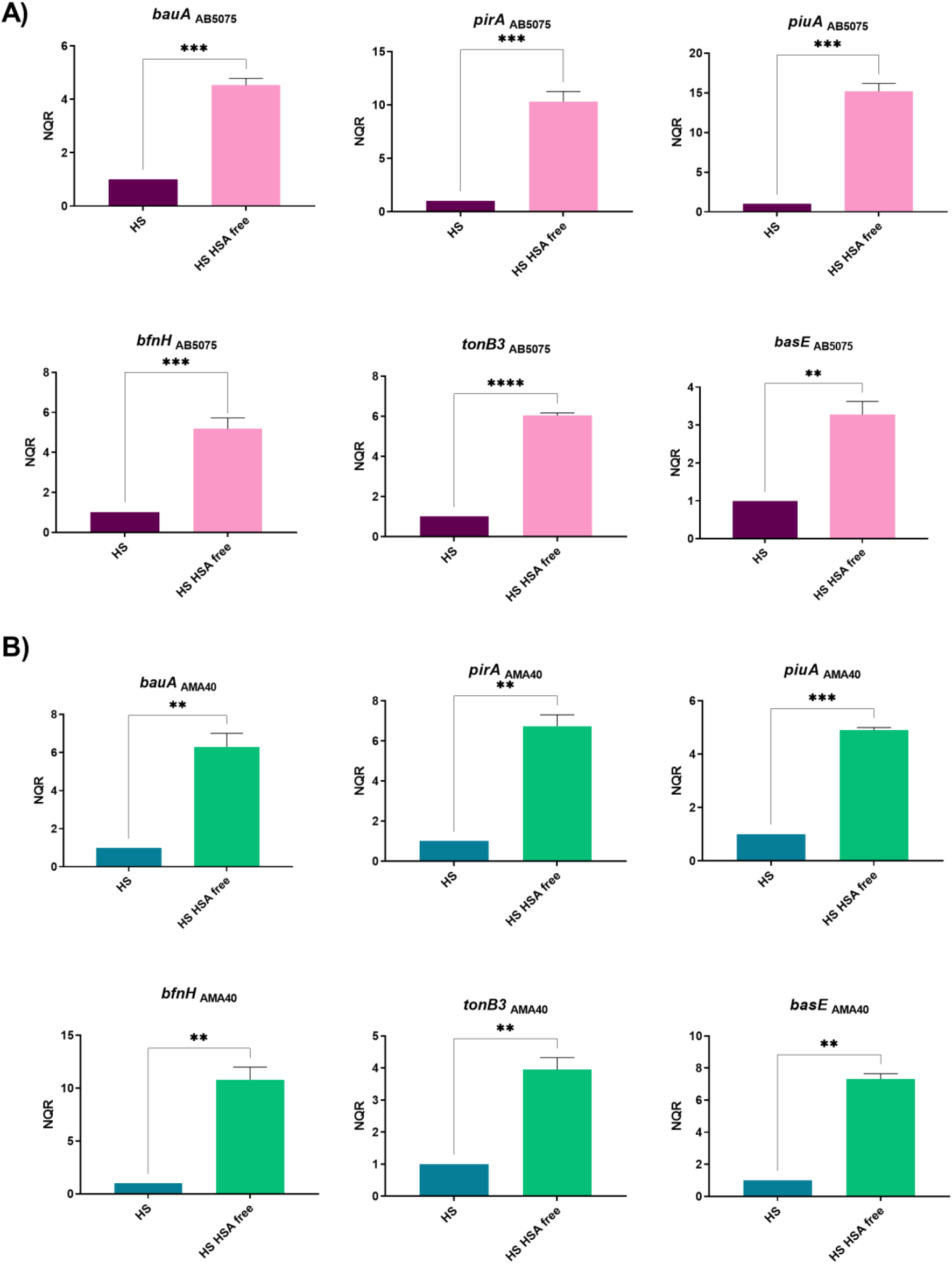
Expression analysis of iron uptake genes in the AB5075 (A) and AMA40 (B) strains. qRT-PCR of the *bauA, pirA, piuA, bfnH, tonB3* and *basE* iron uptake associated genes expressed in the presence of human serum (HS) or HSA-free HS. The data shown are mean ± SD of normalized relative quantities (NRQ) obtained from transcript levels. At least three independent samples were used, and four technical replicates were performed from each sample. The HS condition was used as reference. Statistical significance (*p*< 0.05) was determined by *t* test, one asterisks: *p*< 0.05; two asterisks: *p*< 0.01 and three asterisks: *p*< 0.001.

**Figure 2.**
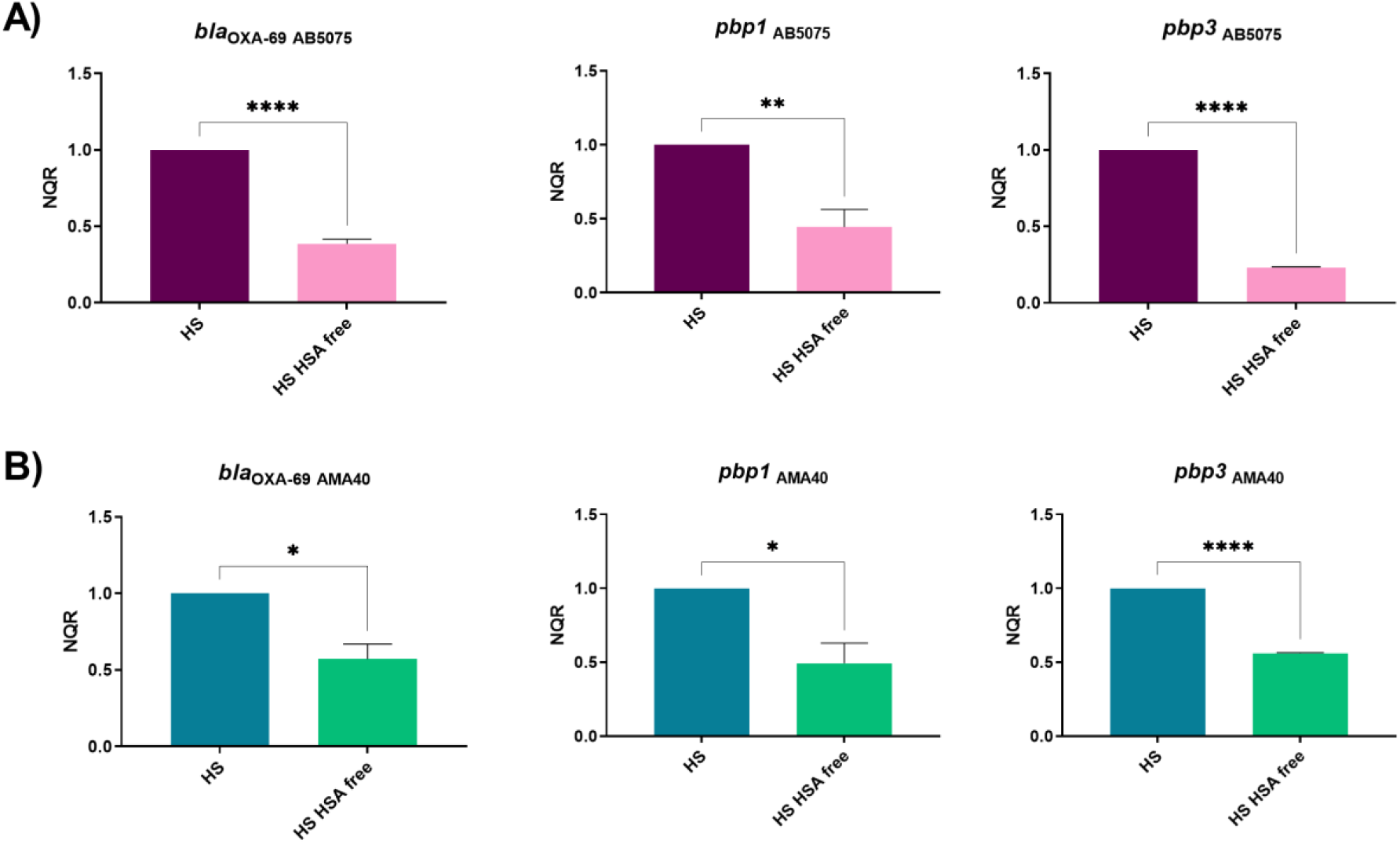
Expression analysis of β-lactamase and PBP genes in the AB5075 (A) and AMA40 (B) strains. qRT-PCR of genes associated with β-lactams resistance, expressed in the presence of human serum (HS) or HSA-free HS. The data shown are mean ± SD of normalized relative quantities (NRQ) obtained from transcript levels. At least three independent samples were used. HS was used as the reference condition. Statistical significance (*p*< 0.05) was determined by *t* test, one asterisks: *p*< 0.05; two asterisks: *p*< 0.01, and three asterisks: *p*< 0.001.

As a genome wide transcriptional response (1120 differentially expressed genes) and an impact in CFDC MICs have been reported when *A. baumannnii* was exposed to HPF (14) (27), we next studied the specific effect of HSA when present in this fluid. Considering that HPF possesses a high HSA-content, but also other components such as reactive oxygen species, monocytes, granulocytes, and other human proteins such the iron chelating protein ferritin, HPF free of HSA was used to assess the differential transcription response of genes coding for iron acquisition functions.

Importantly, qRT-PCR results showed that when the *A. baumannii* AB5075 and AMA40 strains were exposed to HSA free HPF, a statistically significant up-regulation of iron associated genes occurred when compared with untreated HPF (Figures 3A and 3B, Table S1). These results together with those obtained with HS further support the postulated role HSA plays in regulating the transcriptional expression of *A. baumannii* genes coding for transport (*bauA, basE, pirA*, <u>piuA</u> and *bfnH*) and energy transducing (*tonB3*) functions associated with siderophore-mediated iron acquisition processes.

**Figure 3.**
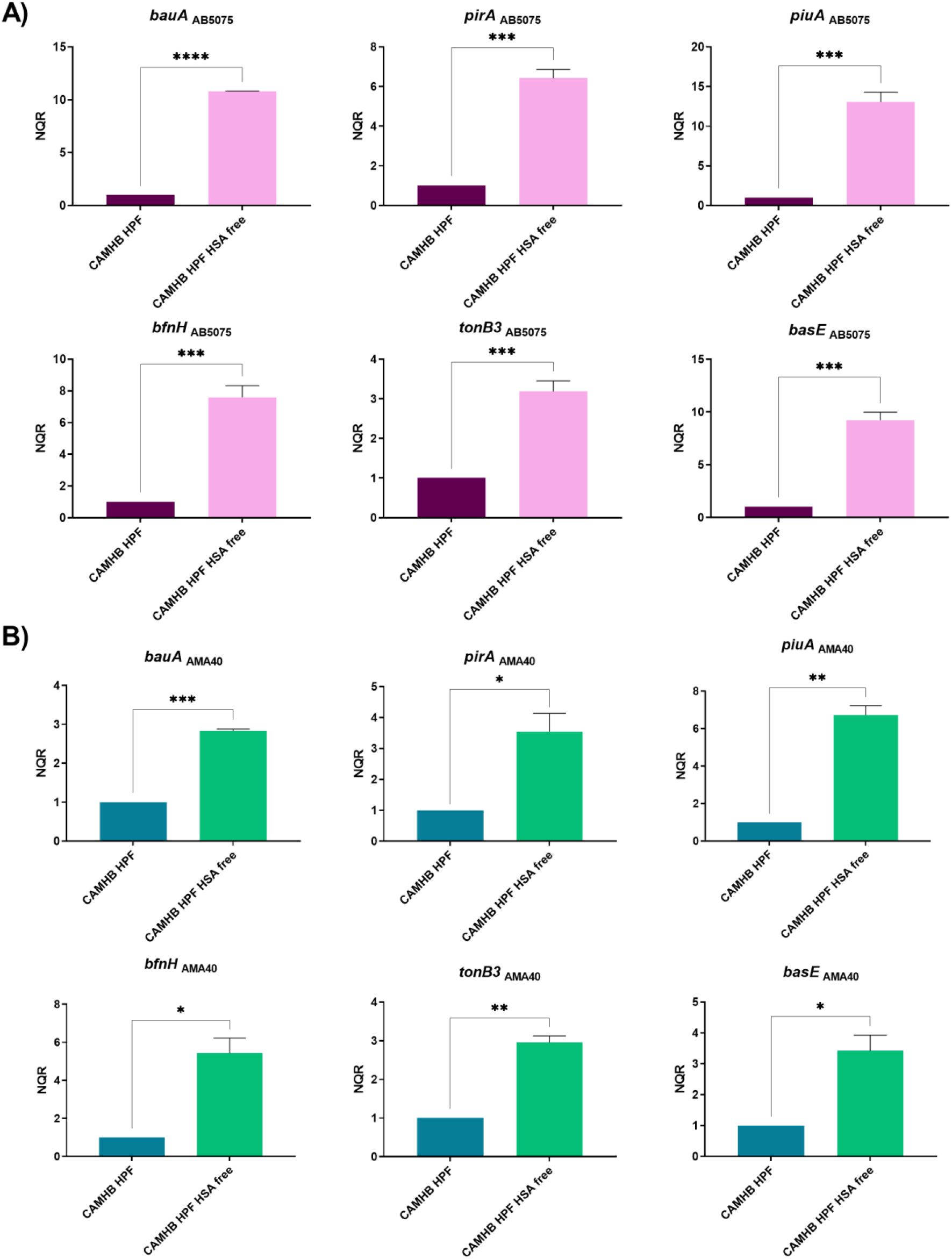
Expression analysis of iron uptake genes in the AB5075 (A) and AMA40 (B) strains. qRT-PCR of the *bauA, pirA, piuA, bfnH, tonB3* and *basE* iron uptake associated genes expressed in cation adjusted Mueller Hinton (CAMHB) supplemented with either human pleural fluid (HPF) or HSA-free HPF. The data shown are mean ± SD of normalized relative quantities (NRQ) obtained from transcript levels. At least three independent samples were used, and four technical replicates were performed from each sample. The CAMHB HPF was used as reference. Statistical significance (*p*< 0.05) was determined by *t* test, one asterisks: *p*< 0.05; two asterisks: *p*< 0.01 and three asterisks: *p*< 0.001.

The transcriptional analysis of the expression of β-lactam resistance genes in HSA -free HPF is aligned with the results seen with HS (Figure 4A and 4B). When *A. baumannii* was cultured in HSA-free HPF, a statistically significant down-regulation of *bla*_OXA-69_ expression was seen for both strains, respectively, as compared with the response displayed by cells cultured in standard HPF (Figures 4A and 4B). In the presence of HSA-free HPF, the expression of *pbp1* was significantly decreased in AB5075 cells cultured in the absence of HSA, while no significant changes were seen for AMA40 (Figures 4A and 4B). A statistically significant down-regulation of *pbp3* expression was seen for AMA40, but not AB5075 when tested using the same experimental conditions (Figure 4B).

**Figure 4.**
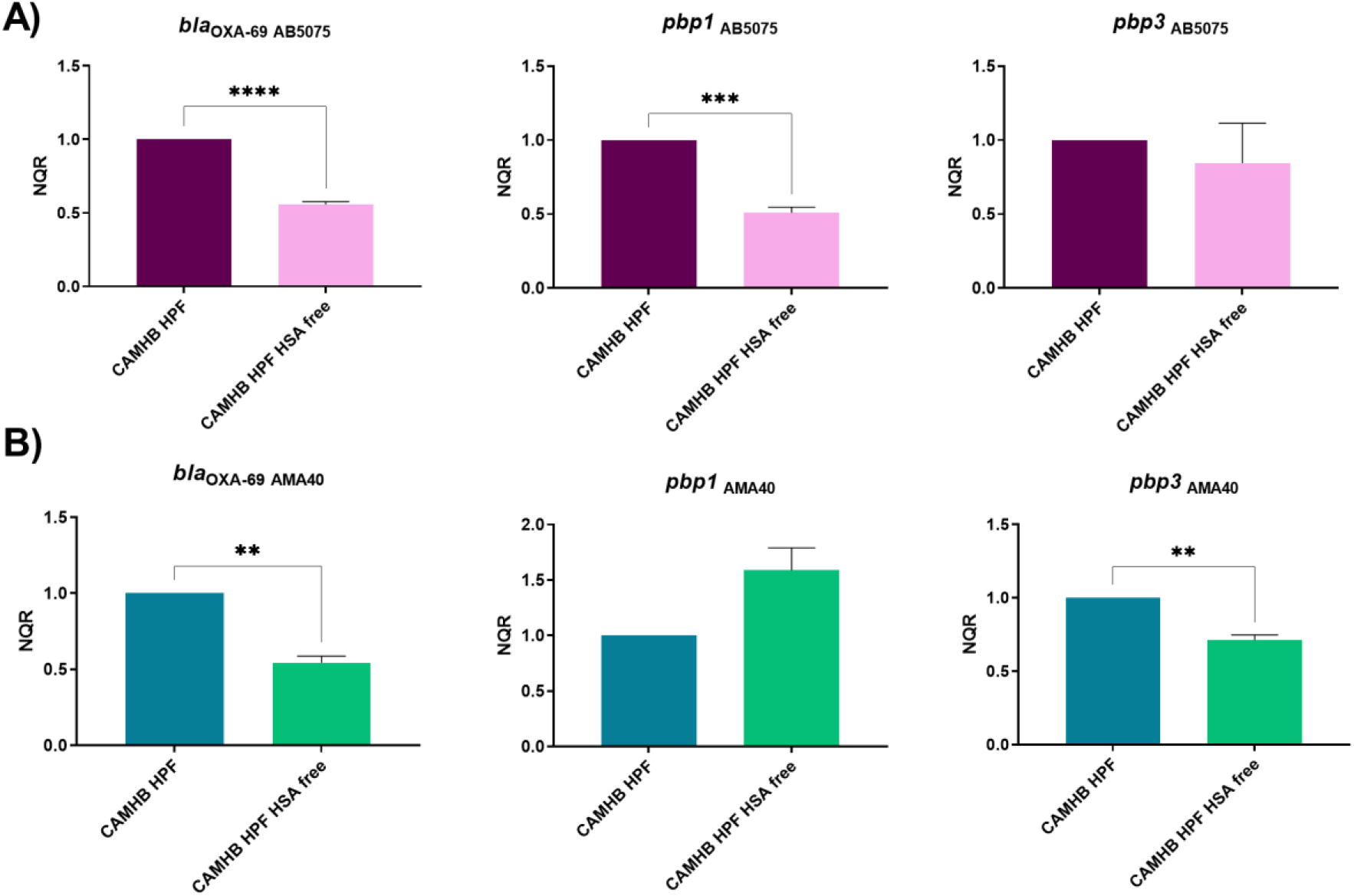
Expression analysis of β-lactamase and PBP genes in the AB5075 (A) and AMA40 (B) strains. qRT-PCR of genes associated with β-lactams resistance expressed in cation adjusted Mueller Hinton (CAMHB) supplemented with either human pleural fluid (HPF) or HSA-free HPF. The data shown are mean ± SD of normalized relative quantities (NRQ) obtained from transcript levels. At least three independent samples were used. CAMHB HPF was used as the reference condition. Statistical significance (*p*< 0.05) was determined by *t* test, one asterisks: *p*< 0.05; two asterisks: *p*< 0.01, and three asterisks: *p*< 0.001.

Taken together, the presented analysis supports the hypothesis that HSA is a significant factor contributing to *A. baumannii* transcriptional responses that have a major impact on CFDC antibacterial potency. In addition, these results further support Le et al. observations that state that fluids with high HSA content (HPF and HS), or pure HSA at a physiological concentration, down-regulate the expression of iron uptake system, genes associated with β-lactam resistance were up-regulated (28). In *A. baumannii*, the role of HSA on affecting the expression of genes involved in its antibiotic resistance and pathogenesis has been previously reported (14, 16, 17, 29, 30). In addition, HSA in combination with carbapenems showed a synergistic increase in natural transformation and expression of competence genes. (28). Ledger et al. showed that HSA directly triggers tolerance to the lipopeptide antibiotic daptomycin in *Staphylococcus aureus* (ref). This tolerance was due to the GraRS two-component system, leading to increased peptidoglycan accumulation increases as well as another independent mechanism that membrane cardiolipin abundance (25). Notably, these investigators also showed the specific and direct role of HSA as the molecule leading the observed effects with *S. aureus* (25). In addition, the role of HSA on augmenting the virulence was not only seen in bacteria. In pathogenic fungi, such as *Candida glabrata*, the presence of HSA also contributes to the virulence of this species (26).

### Role of ferric HSA on *A. baumannii* response affecting CFDC susceptibility

The results obtained with HSA-free fluids strongly support the possibility that HSA is the molecule modulating changes in the expression of genes coding for iron acquisition and β-lactam resistance functions, both of which contribute to alterations in CFDC MICs. However, the mechanisms by which HSA triggers these effects is not yet fully understood. HSA can be playing at least two different possible roles: *i*) HSA exerts a direct role in the modulation of virulence-associated phenotypes ; or ii) serves as a carrier of metal ions, such as iron (31, 32), affecting the differential expression of genes coding for active iron uptake systems and β-lactam resistance genes, and as consequence affecting CFDC’s efficacy.

With the aim of evaluating if HSA is acting as an iron carrier causing the down-regulation of genes associated with iron-uptake ultimately affecting CFDC activity, iron was removed from HSA (Fe-free HSA). The transcriptional analysis by qRT-PCR showed that when *A. baumannii* was exposed to Fe-free HSA, a statistically significant up-regulation of iron acquisition associated genes occurred in both strains evaluated (Figures 5A and 5B). In addition, when Fe-free HSA was supplemented with FeCl_3_ or untreated HSA, the transcription expression levels of the tested genes were restored to levels comparable to those of samples incubated with untreated HSA. These results indicate that the iron carried by HSA plays a role in regulating the expression of *A. baumannii* iron-acquisition genes.

**Figure 5.**
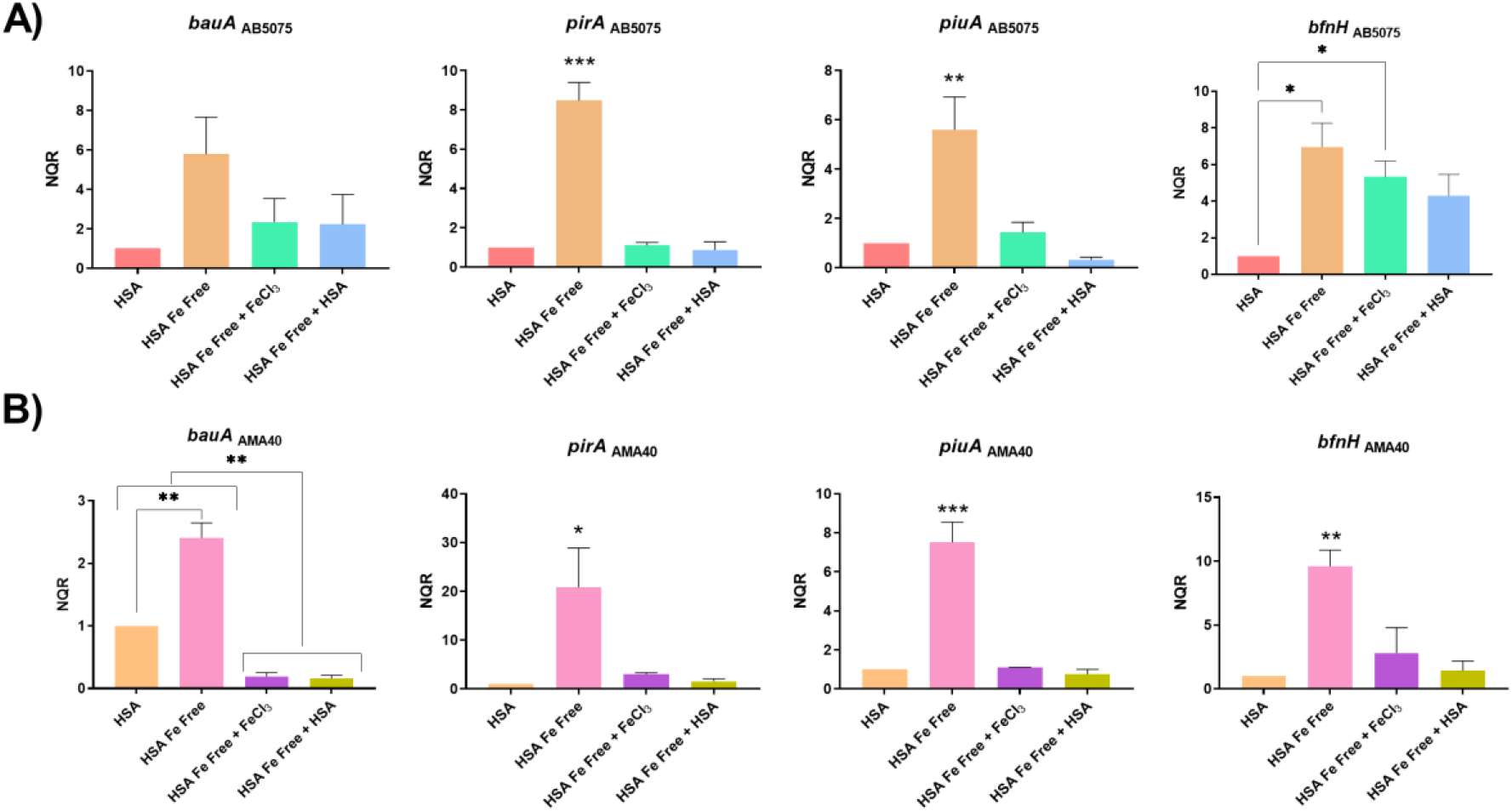
Genetic analysis of iron uptake genes of AB5075 (A) and AMA40 (B) *A. baumannii* strains. qRT-PCR of genes associated with iron uptake, *bauA, pirA, piuA, tonB3*, and *bfnH* expressed in human serum albumin (HSA), HSA Fe free, HSA Fe free supplemented with FeCl_3_ or HSA Fe free supplemented with HSA. The data shown are mean ± SD of normalized relative quantities (NRQ) obtained from transcript levels. At least three independent samples were used, and four technical replicates were performed from each sample. The HSA condition was used as reference. Statistical significance (*p*< 0.05) was determined by ANOVA followed by Tukey’s multiple-comparison test, one asterisks: *p*< 0.05; two asterisks: *p*< 0.01 and three asterisks: *p*< 0.001.

Expression levels of β-lactam resistance genes in Fe-free HSA were hext evaluated (Figures 6A and 6B). *A. baumannii* cells cultured in Fe-free HSA, showed a decreased in *bla*_OXA-69_, *pbp1*, and *pbp3* transcripts for both strains (Figures 6A and 6B) with respect to the untreated HSA condition. However, when Fe-free HSA was supplemented with FeCl_3_ or HSA, the expression levels were restored to those of the HSA condition (Figures 6A and 6B). Statistically significantly increases varied depending on the evaluated gene, condition, and strain (Figures 6A and 6B).

**Figure 6.**
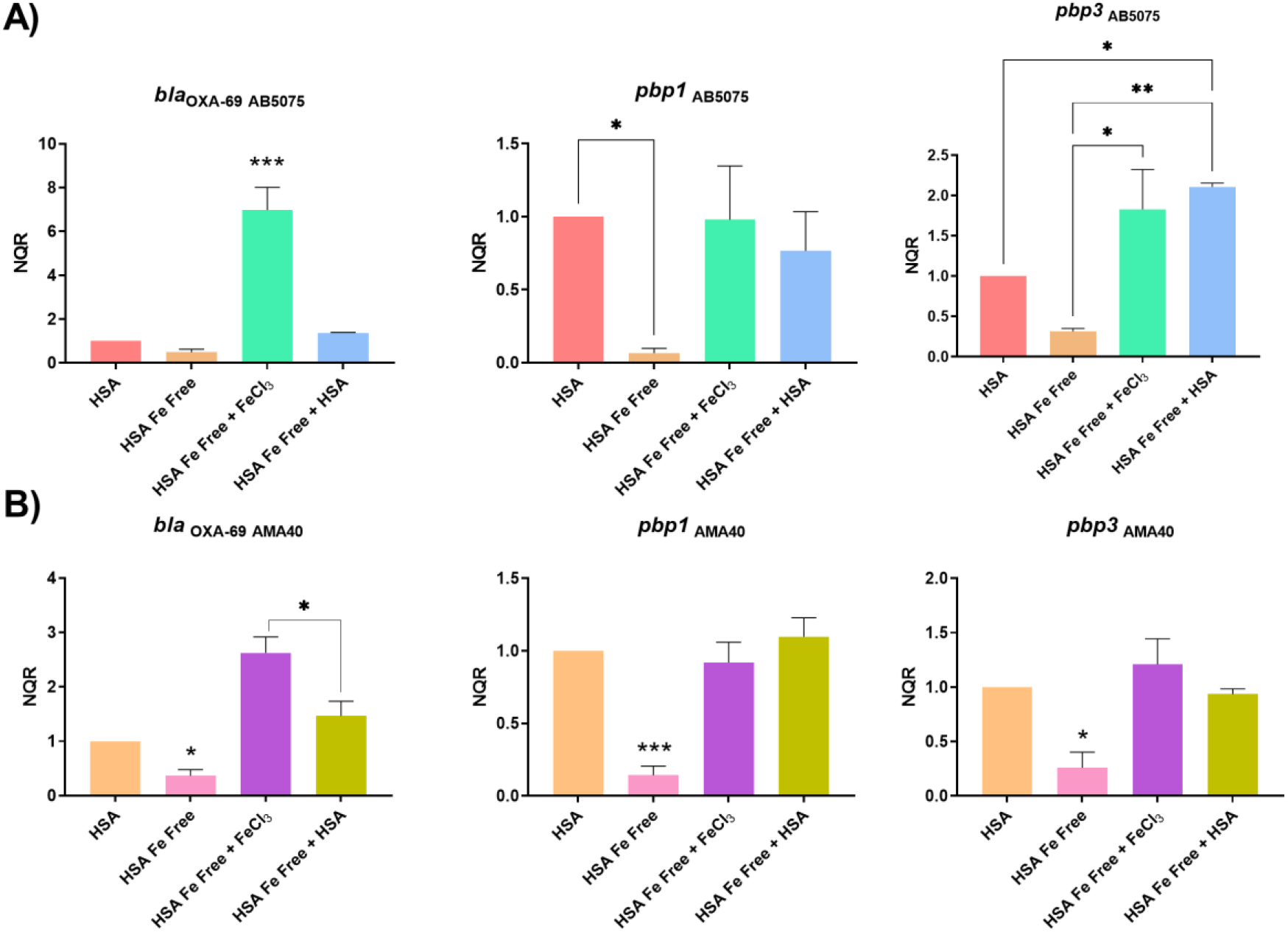
Genetic analysis of β-lactamase and PBP genes of AB5075 (A) and AMA40 (B) *A. baumannii* strains. qRT-PCR of genes associated with β-lactams resistance expressed in human serum albumin (HSA), HSA Fe free, HSA Fe free supplemented with FeCl_3_ or HSA Fe free supplemented with HSA. The data shown are mean ± SD of normalized relative quantities (NRQ) obtained from transcript levels. At least three independent samples were used. HSA was used as the reference condition. Statistical significance (*p*< 0.05) was determined by ANOVA followed by Tukey’s multiple-comparison test, one asterisks: *p*< 0.05; two asterisks: *p*< 0.01, and three asterisks: *p*< 0.001.

Studies *in vitro* have shown that the intrinsic activity of CFDC against *Pseudomonas aeruginosa* is enhanced under iron-limited conditions (9), showing that supplementation with ferric iron increases CFDC MICs. To assess if the same effect occurs in *A. baumannii*, we tested the role of free iron on AMA50 and AB5075 CFDC MICs *in vitro*. We observed that a significant increase in CFDC MICs was noted when the iron depleted media (CAMHA) was supplemented with 20 μM FeCl_3_ or 40 μM FeCl_3_ (Table S2). This results not only shows that the effect of iron on CFDC is not pathogen specific, but also suggests that the variations in the iron content of different human fluids (free iron or iron bind to human proteins) could be a potential factor that affects the efficacy of CFDC.

We next decided to determine the effect of ferric HSA at the phenotypic level through changes on the susceptibility of the bacteria to CFDC. CFDC MICs for AMA40 and AB5075 using CAMHB (untreated), CAMHB supplemented with HSA pre-Chelex® treatment (HSA Fe) or Fe-Free HSA were performed. In addition, the CAMHB was supplemented with Fe-Free HSA + 100μM FeCl3 and Fe-Free HSA + 3.5% HSA to further support the observation. The Minimal bactericidal concentration (MBC) of the mentioned conditions was also performed keeping in mind the occurrence of heteroresistance that cannot be detected using the microdilution method. A decreased in the MIC and MBC values for AB5075 were observed when the iron was removed from HSA, values that were restored or even increased more when inorganic iron was added back to the Fe-free HSA tested condition (Table 2). A similar response was observed when untreated HSA was added to the medium. For the AMA40 strain, the same effect on the CFDC MIC values was observed (Table 2). The MIC and MBC values showing a decreased when iron was removed, while restored or increased values were observed when iron or HSA were added (Table 2). These results indicate that iron-rich HSA and/or the presence of inorganic iron are associated with reduced susceptibility to CFDC.

**Table 2:**
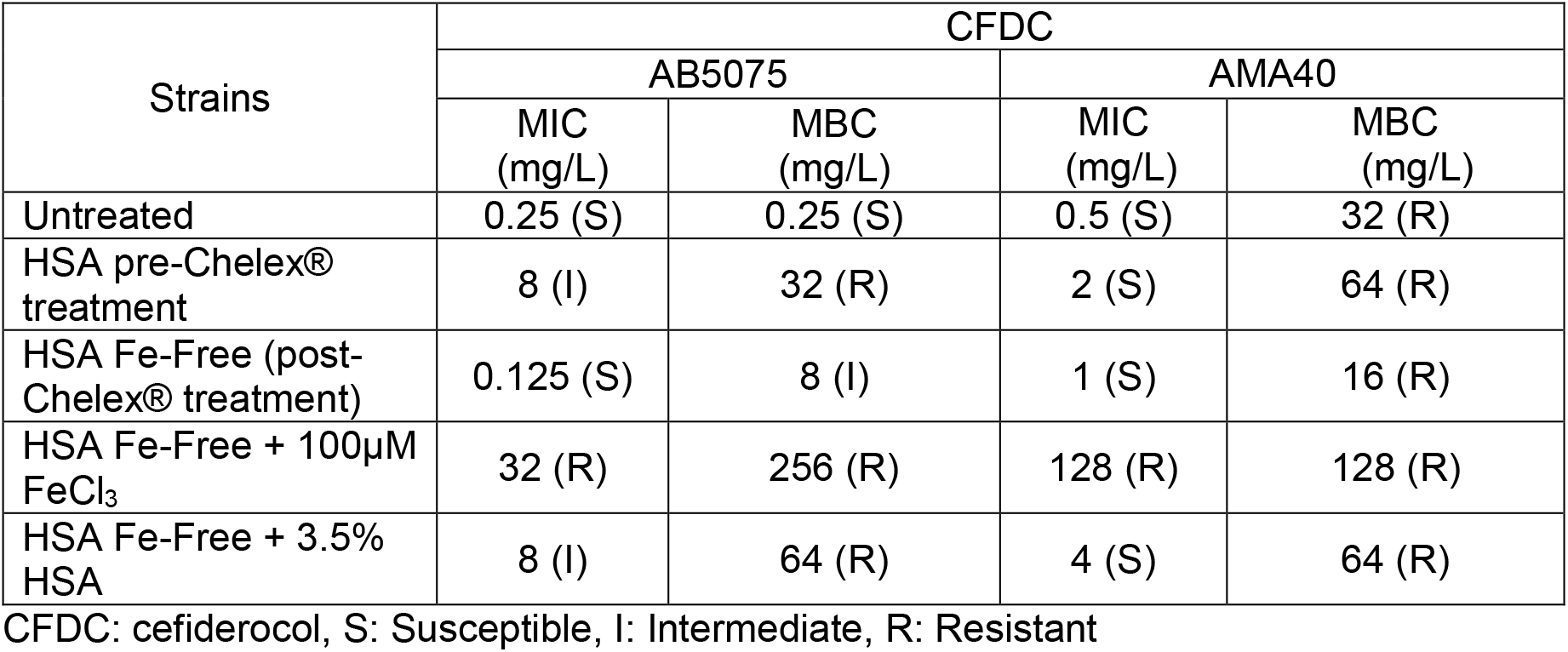
Minimal Inhibitory Concentrations (MICs) and Minimal Bactericidal Concentration (MBCs) of CFDC for the CRAB AB5075 and AMA40 strains, performed by microdilution in iron-depleted CAMHB and CAMHB with different experimental conditions.

| Strains | CFDC |  |  |  |
| --- | --- | --- | --- | --- |
|  | AB5075 |  | AMA40 |  |
|  | MIC (mg/L) | MBC (mg/L) | MIC (mg/L) | MBC (mg/L) |
| Untreated | 0.25 (S) | 0.25 (S) | 0.5 (S) | 32 (R) |
| HSA pre-Chelex® treatment | 8 (I) | 32 (R) | 2 (S) | 64 (R) |
| HSA Fe-Free (post-Chelex® treatment) | 0.125 (S) | 8 (I) | 1 (S) | 16 (R) |
| HSA Fe-Free + 100µM FeCl<sub>3</sub> | 32 (R) | 256 (R) | 128 (R) | 128 (R) |
| HSA Fe-Free + 3.5% HSA | 8 (I) | 64 (R) | 4 (S) | 64 (R) |

CFDC: cefiderocol, S: Susceptible, I: Intermediate, R: Resistant

Considering these observations, HSA can serve as a host heme-scavenging protein as seen in other gram-positive bacteria such as *Staphylococcus aureus, Streptococcus dysgalactiae* and *Streptococcus mutans* (23, 25). HSA can exert a direct role on the transcriptional and phenotypical response of *A. baumannii*. Pinsky *et al*., showed that HSA facilities the iron utilization by *C. albicans* (22). In addition, *C. glabrata* interact with albumin while colonizing and infecting the vaginal mucosa. These authors concluded that albumin helps *C. glabrata* to overcome iron restriction (26). However, as mentioned above, there are evidence that suggested a different role (33). For example, in *P. aeruginosa*, in the early exponential phase, HSA present in serum increased the expression of iron-related genes in an iron-independent manner (33).

### Changes in the expression of genes coding for iron uptake functions and β-lactam resistance in cerebrospinal fluid (CSF), a low HSA content fluid

Previous work showed that a human fluid, such as urine, with none or traces of HSA triggers a different behavioral and transcriptional response by *A. baumannii* (27). This work also showed that CFDC MICs values are not significantly modified, while the expression of *piuA, pirA, bauA*, and *bfnh* was enhanced when bacteria were cultured in urine, suggesting that CFDC uptake through active iron transport systems is not impaired (27).

Since, the incidence of *A. baumannii* as a causal agent of nosocomial meningitis has been increasing in the last years with a mortality rate of 15-71% (34, 35), and knowing that only 1% of the cerebrospinal fluid (CSF) content corresponds to proteins, where HSA represents 70% of these proteins (36), we decided to study the expression of genes coding for iron uptake systems and β-lactams resistance when *A. baumannii* cells are exposed to 20% CSF. It is important to note that our analysis resulted in no detectable iron in 100% and 20% CSF samples (see Table S3). qRT-PCR showed that exposure of *A. baumannii* to CSF resulted in a statistically significant up-regulation of *piuA, pirA*, and *bfnH* with both tested strains when compared to the CAMHB control (Figures 7A and 7B). In contrast, *bauA* transcription was down-regulated when compared to the same control (Figures 7A and 7B). In addition, as we have used 4% CSF before when performing the RNA-seq analysis (15), we decided to conduct the same analysis (Figure S1) observing similar results to those collected using 20% CSF. These observations agree with Nishimura et al. results, showing a similar differential expression of iron acquisition genes when *A. baumannii* encounters low HSA content human fluids (27). Moreover, the expression levels of β-lactam resistance genes are also affected by the presence of CSF; while it promotes the down-regulation of *bla*_OXA-69_, it up-regulates *pbp1*, and *pbp3* transcription in AB5075 and AMA40 (Figures 8A and 8B) at both CSF concentrations (Figure S2). As expected, considering the previous results seen in urine, changes in the MIC to CFDC were not observed when *A. baumannii* AB5075 or AMA40 were exposed to CSF (Table S3).

**Figure 7.**
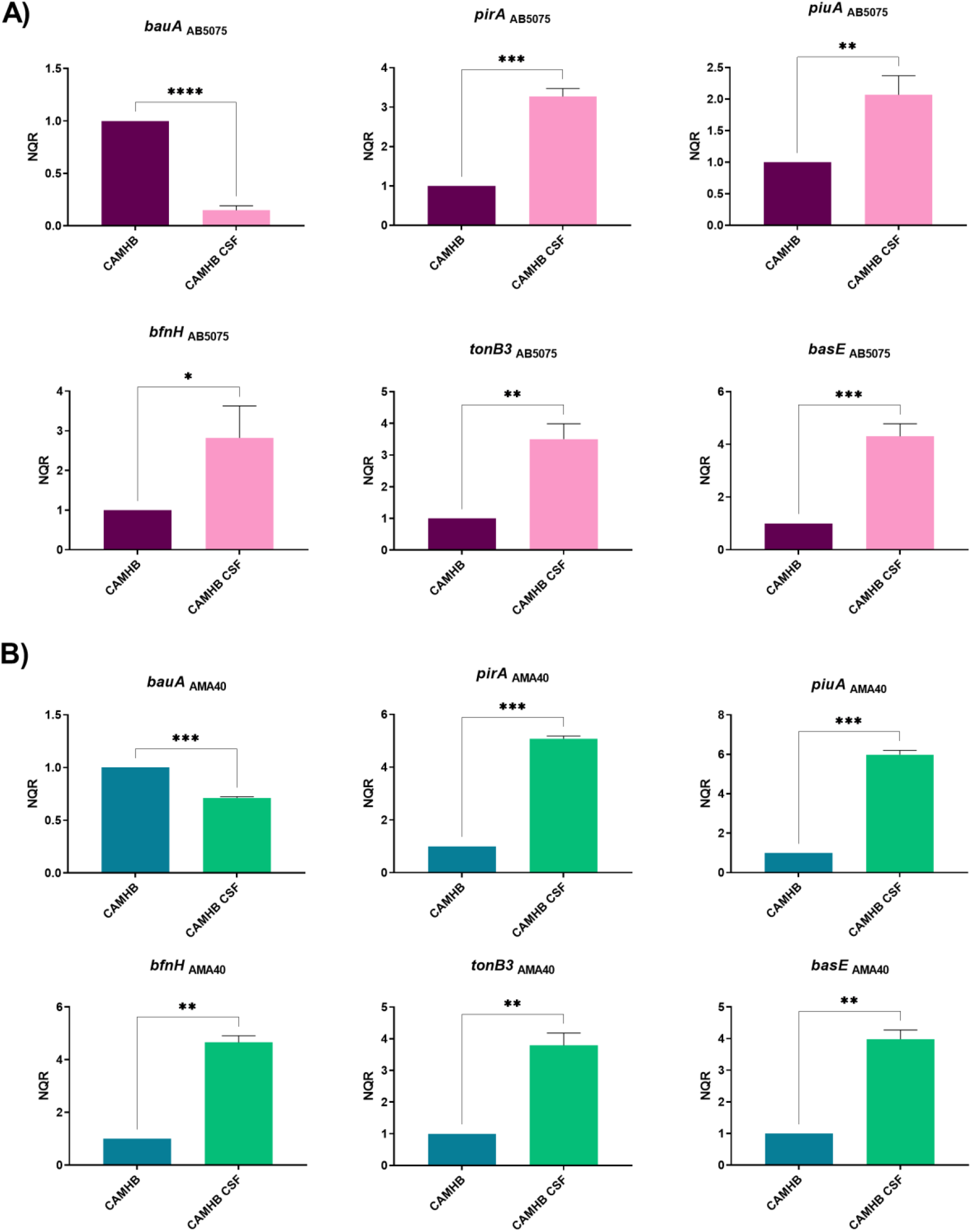
Genetic analysis of iron uptake genes of AB5075 (A) and AMA40 (B) *A. baumannii* strains. qRT-PCR of genes associated with iron uptake, *bauA, pirA, piuA, bfnH, tonB3*, and *basE* expressed in cation adjusted Mueller Hinton (CAMHB) or CAMHB supplemented with cerebrospinal fluid (CSF). The data shown are mean ± SD of normalized relative quantities (NRQ) obtained from transcript levels. At least three independent samples were used, and four technical replicates were performed from each sample. The CAMHB was used as reference. Statistical significance (*p*< 0.05) was determined by *t* test, one asterisks: *p*< 0.05; two asterisks: *p*< 0.01, and three asterisks: *p*< 0.001.

**Figure 8.**
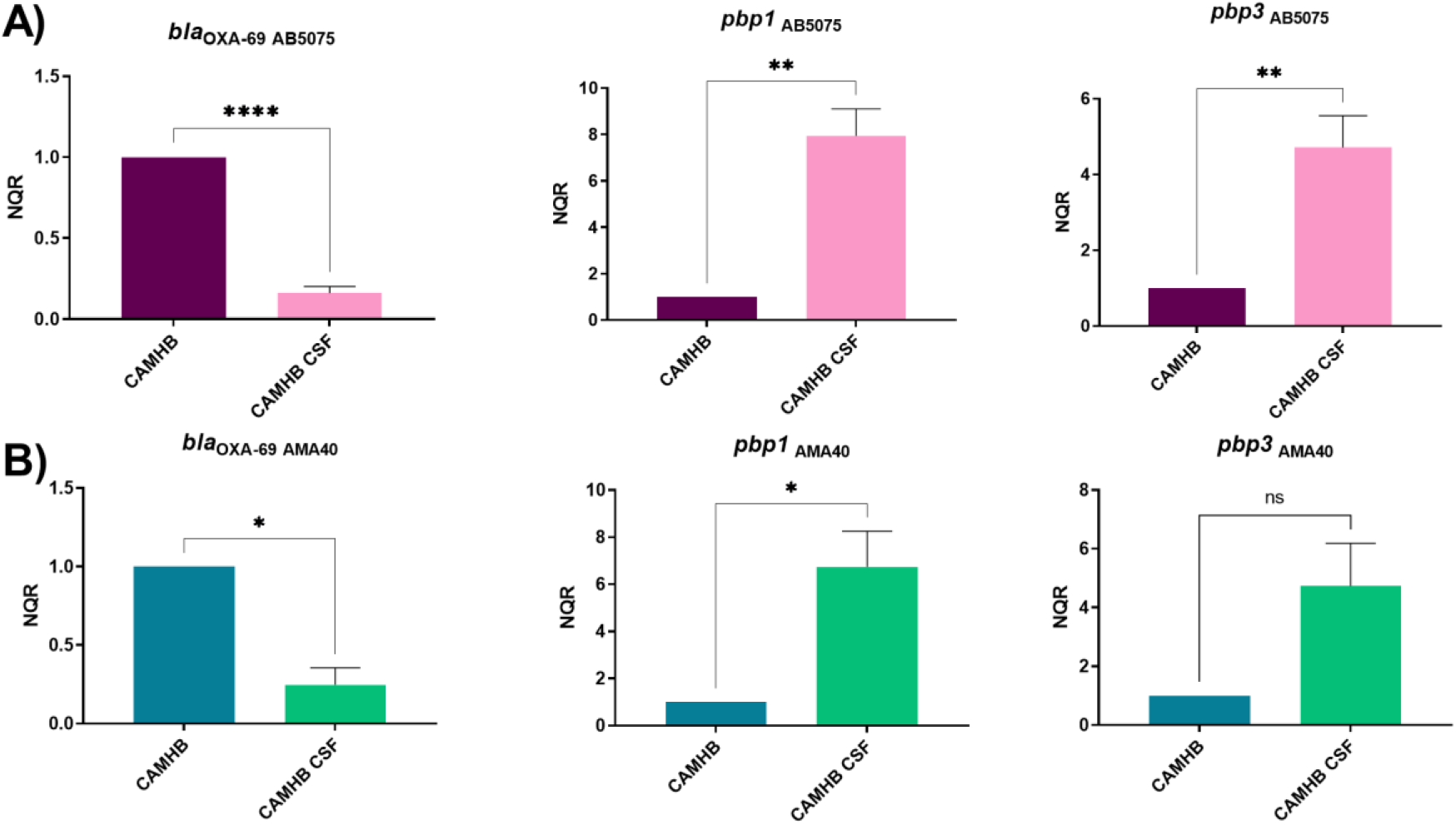
Genetic analysis of β-lactamase and PBP genes of AB5075 (A) and AMA40 (B) *A. baumannii* strains. qRT-PCR of genes associated with β-lactams resistance, expressed in cation adjusted Mueller Hinton (CAMHB) or CAMHB supplemented with cerebrospinal fluid (CSF). The data shown are mean ± SD of normalized relative quantities (NRQ) obtained from transcript levels. At least three independent samples were used. LB was used as the reference condition. Statistical significance (*p*< 0.05) was determined was determined by *t* test, one asterisks: *p*< 0.05; two asterisks: *p*< 0.01, and three asterisks: *p*< 0.001.

In summary, these results support our hypothesis that the absence of detectable iron and the low HSA content of CSF compared to HS or HPF are the signals that affect the antimicrobial activity of CFDC.

## CONCLUDING REMARKS

The results obtain in the present work further expand previous investigations and provide insights into earlier observations suggesting that high HSA containing human fluids, including HPF and HS, affect the potency of CFDC as an antibacterial agent against *A. baumannii*. Here we demonstrate that HSA is the molecule involved in inducing the changes at the expression of genes that facilitate iron transport and increases expression of β-lactamase associated genes leading to increases in CFDC MICs. The removal of iron from HSA suggests that the ability of HSA to bind iron and provide an iron-rich environment play a role by reducing the expression of genes coding for siderophore-mediated iron acquisition that are critical for the uptake and the ensuing antimicrobial activity of catechol-antibiotic conjugates such as CFDC. On the other hand, the mechanism by which the presence of ferric HSA controls the differential expression of genes coding for β-lactam resistance remains to be elucidated.

## MATERIALS AND METHODS

### Bacterial strains

The carbapenem-resistant *A. baumannii* AB5075 (*bla*_OXA-23_ and *bla*_OXA-51_) (28, 30, 37) model strain and the clinical carbapenem-resistant AMA40 (*bla*_NDM-1_ and *bla*_OXA-51_) isolate (27, 38, 39) were used in this work.

### RNA Extraction, Quantitative Reverse Transcription Polymerase Chain Reaction (qRT-PCR)

*A. baumannii* AB5075 and AMA40 cells were cultured in iron depleted cation adjusted Mueller Hinton (CAMHB) and incubated with agitation for 18 h at 37°C. Overnight cultures were then diluted 1:10 in fresh CAMHB, CAMHB supplemented with 3.5 % human serum albumin (HSA), 3.5 % iron-free HSA, 3.5 iron-free HSA + 100 μM FeCl_3_, 3.5 % iron-free HSA + 100 μM FeCl_3_ + 3.5 % HSA, or CAMHB supplemented with 4 % human pleural fluid (HPF), CAMHB supplemented with HSA-free HPF, 100 % human serum (HS) or 100 % HSA free HS. All samples were incubated with agitation for 18 h at 37°C. RNA was immediately extracted using the Direct-zol RNA Kit (Zymo Research, Irvine, CA, USA) following manufacturer’s instructions as previously described (27). Total RNA extractions were performed in three biological replicates for each condition. The extracted and DNase treated RNA was used to synthesize cDNA iScriptTM Reverse Transcription Supermix for qPCR (Bio-Rad, Hercules, CA, USA) using the manufacturer’s protocol. The cDNA concentrations were adjusted to 50 ng/μL and qPCR was conducted using the qPCRBIO SyGreen Blue Mix Lo-ROX following manufacturer’s protocol (PCR Biosystems, Wayne, PA, USA). At least three biological replicates of cDNA were each tested in triplicate using the CFX96 TouchTM Real-Time PCR Detection System (Bio-Rad, Hercules, CA, USA). Data are presented as NRQ (Normalized relative quantities) calculated by the qBASE method (40), using *recA* and *rpoB* genes as normalizers. Asterisks indicate significant differences as determined by *t* test or ANOVA followed by Tukey’s multiple comparison test (*p* < 0.05), using GraphPad Prism (GraphPad software, San Diego, CA, USA).

### HSA removal

To remove the HSA from HS or HPF, the ProteoExtract® Albumin/IgG Removal Kit (Sigma-Aldrich, MA, United States) was used following the manufacturer’s instructions. To corroborate the correct removal of HSA, protein samples of both fluid pre and post HSA-removal-treatment were separated by SDS-PAGE and visualized by Coomassie staining (Figure S3). Protein concentrations in soluble extracts were determined by the Bradford method (41) pre and post HSA-removal-treatment (Table S4).

### Iron removal

Iron was removed from has samples using Chelex® 100 Chelating Ion Exchange Resin (Bio-Rad, Hercules, CA, USA) following the manufacturer’s instructions. The iron content of pre and post Chelex® 100 treatment was determined using the Iron Assay Kit (Sigma-Aldrich, MA, United States) following the manufacturer’s recommendations (Table S5).

### Antimicrobial Susceptibility Testing

Antibiotic susceptibility assays were performed following the procedures recommended by the Clinical and Laboratory Standards Institute (CLSI). After OD_600_ adjustment, 100 μL of *A. baumannii* AB5075 and AMA40 cells grown in fresh CAMHB, or CAMHB supplemented with human fluids (HPF, HS or CSF) or fluids where HSA was removed (HSA-free HPF, and HSA-free HS) were inoculated on CAMH agar plates (CAMHA) as previously described (27). Antimicrobial commercial E-strips (Liofilchem S.r.l., Roseto degli Abruzzi, Italy) CFDC were used. CAMHA plates were incubated at 37ºC for 18 h. CLSI breakpoints were used for interpretation (42). *E. coli* ATCC 25922 was used for quality control purposes.

The Microdilution test was used to study the effect of HSA and iron-free HSA on CFDC MICs. CAMHB was prepared as described above and supplemented with HSA, iron-free HSA, iron-free HSA Fe + 100μM FeCl_3_ or iron-free HSA + iron-free HSA Fe _ 3.5% HSA to performed CFDC (range 0.25-512 mg/L) following the CLSI guidelines (42). The Quality control strain (*Escherichia coli* ATCC 25922) was tested for as control purposes (42, 43). CLSI breakpoints were used for interpretation (42).

To study the role of ferric iron on AMA40 and AB5075 CFDC MIC, iron-depleted CAMHA and iron-depleted CAMHA supplemented with 20 μM FeCl_3_ or 40 μM FeCl_3_ were used. The MICs were determine using CFDC MTS strips (Liofilchem S.r.l., Italy) following the CLSI guidelines (42).

## Author Contributions

B.N., J.E., M.R.T., T.S., C.P., N.G., V.M., F.P., C.R., R.A.B, M.E.T. and M.S.R. conceived the study and designed the experiments. B.N., J.E., M.R.T., T.S., C.P., N.G., V.M., F.P., C.R., R.S. and M.S.R. performed the experiments and genomics and bioinformatics analyses. M.R.T., T.S., F.P., R.A.B, M.E.T. and M.S.R. analyzed the data and interpreted the results. R.A.B., M.E.T. and M.S.R. contributed reagents/materials/analysis tools. M.R.T., T.S., F.P., R.A.B, M.E.T. and M.S.R. wrote and revised the manuscript. All authors read and approved the final manuscript.

## Supplementary material

**Table S1:** Comparison of the level of expression of iron associated and antibiotic resistance genes obtained by qRT-PCR in the two CRAB strains.

| Gene name | AB5075 (Log<sub>2</sub>FC) |  | AMA40 (Log<sub>2</sub>FC) |  |
| --- | --- | --- | --- | --- |
|  | HS HSA free | HPF HSA free | HS HSA free | HPF HSA free |
| **Iron Uptake** |  |  |  |  |
| *bauA* | 2,18 | 3,44 | 2,65 | 1,50 |
| *pirA* | 3,36 | 2,68 | 2,75 | 1,82 |
| *piuA* | 3,92 | 3,70 | 2,29 | 2,75 |
| *bfnH* | 2,37 | 2,92 | 3,43 | 2,43 |
| *tonB3* | 2,59 | 1,67 | 1,98 | 1,56 |
| *basE* | 1,71 | 3,20 | 2,87 | 1,77 |
| **Antibiotic resistance** |  |  |  |  |
| *bla<sub>OXA-69</sub>* | -1,38 | -0,84 | -0,82 | -0,88 |
| *pbp1* | -1,20 | -0,97 | -1,05 | 0,66 |
| *pbp3* | -2,10 | -0,28 | -0,84 | -0,49 |

- Log<sub>2</sub>FC > 1 (p<0.05) ● Log<sub>2</sub>FC < - 1 (p<0.05) - Log<sub>2</sub>FC (0-1) ● Log<sub>2</sub>FC (-1-0)

**Table S2:** Minimal Inhibitory Concentrations (MICs) of cefiderocol (CFDC) for the CRAB AB5075 and AMA40 strains performed using CFDC MTS strips (Liofilchem S.r.l., Italy) on Iron-depleted CAMHA (Cation Adjusted Mueller Hinton Agar) supplemented with 20 μM or 40 μM FeCl_3_.

| Strain | CAMHA | CAMHA + 20 μM FeCl<sub>3</sub> | CAMHA + 40 μM FeCl<sub>3</sub> |
| --- | --- | --- | --- |
| AMA40 | 0.25 | 1.5 | 3.0 |
| AB5075 | 0.19 | 1.5 | 2.0 |

CAMHA: Iron-depleted Cation Adjusted Mueller Hinton (treated with CHELEX as recommend by the CLSI).

**Table S3:** Minimal Inhibitory Concentrations (MICs) of cefiderocol (CFDC) for the CRAB AB5075 and AMA40 strains, performed using CFDC MTS strips (Liofilchem S.r.l., Italy) on Iron-depleted CAMHA (Cation Adjusted Mueller Hinton Agar) and supplemented with 4% or 20% of cerebrospinal fluid (CSF).

| Condition | CFDC MIC (*mg/L*) |  |
| --- | --- | --- |
|  | AB5075 | AMA40 |
| CAMHB | 0.50 (S) | 0.50 (S) |
| 4% CSF | 0.50 (S) | 0.50 (S) |
| 20% CSF | 0.38 (S) | 0.75 (S) |

CFDC: cefiderocol, S: Susceptible, R: Resistant

**Table S4.** Total protein concentration determined in CAMHB, and different fluids analyzed using the ProteoExtract® Albumin/IgG Removal Kit (Sigma-Aldrich, MA, United States).

|  | Protein concentration (mg/mL) | Reference |
| --- | --- | --- |
| CAMHB | BLQ | This work |
| HPF | 22.66 ± 4.53 | This work |
| HPF HSA Free | BLQ | This work |
| HS | 14.15 ± 0.31 | This work |
| HS HSA Free | BLQ | This work |

CAMHB: iron depleted cation adjusted Mueller Hinton, HSA: human serum albumin, HPF: human pleural fluids, HS: human serum, CSF: cerebrospinal fluid, BLQ: below limit of quantification.

**Table S5.** Total Iron concentration determined in CAMHB, and different fluids analyzed using the Iron Assay Kit (Sigma-Aldrich, MA, United States).

|  | [Fe] total (μM) | Reference |
| --- | --- | --- |
| CAMHB | BLQ | This work |
| HSA | 65.4 ± 0.39 | This work |
| HSA Fe Free | BLQ | This work |
| HPF | 129.97 ± 0.86 | [1] |
| HPF HSA Free | BLQ | This work |
| HS | 65.45 ± 0.46 | This work |
| HS HSA Free | BLQ | This work |
| CSF | BLQ | This work |

CAMHB: iron depleted cation adjusted Mueller Hinton, HSA: human serum albumin, HPF: human pleural fluids, HS: human serum, CSF: cerebrospinal fluid, BLQ: below limit of quantification.

---

**Figure S1.**
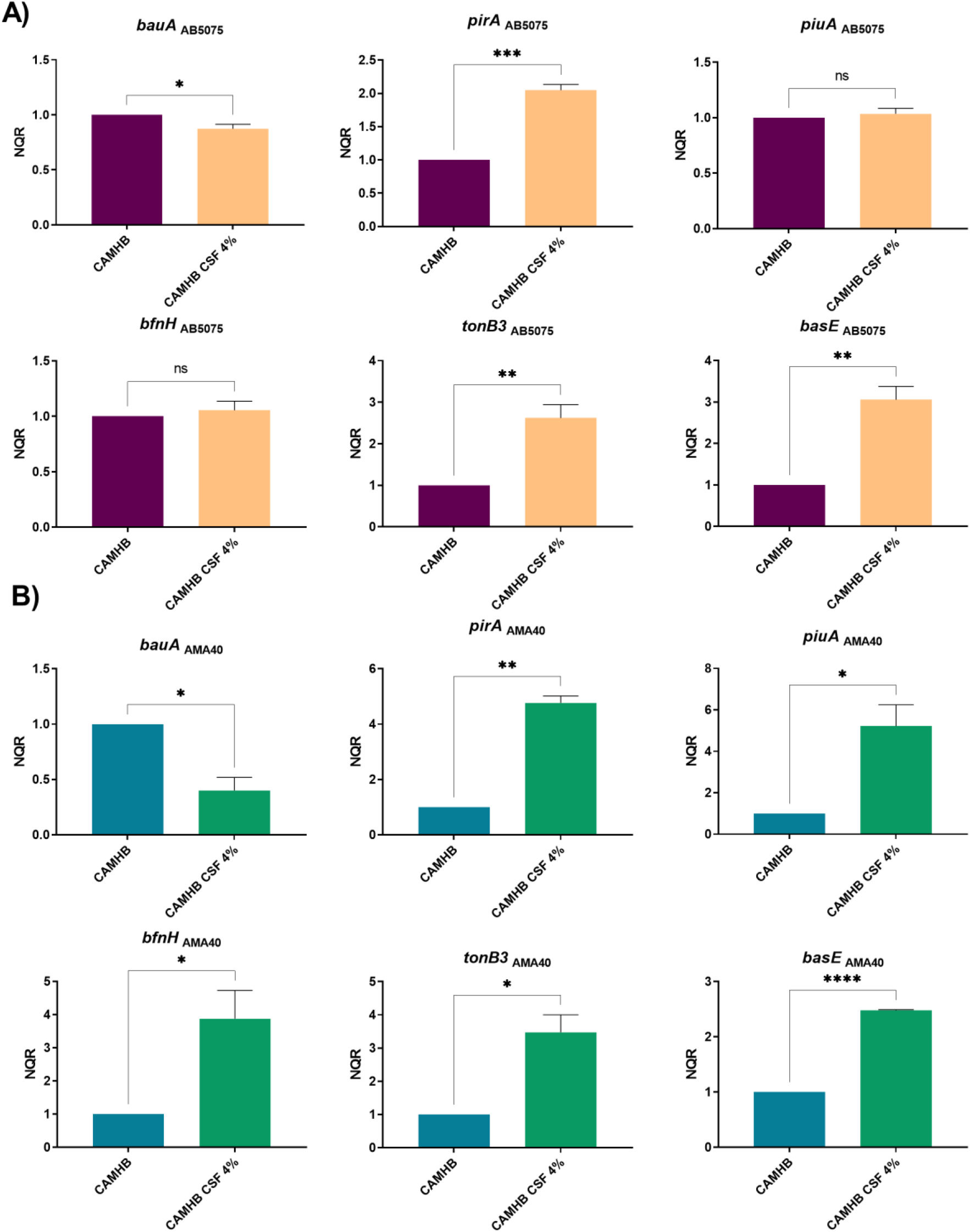
Genetic analysis of iron uptake genes of AB5075 (A) and AMA40 (B) *A. baumannii* strains. qRT-PCR of genes associated with iron uptake, *bauA, pirA, piuA, bfnH, tonB3*, and *basE* expressed in cation adjusted Mueller Hinton (CAMHB) or CAMHB supplemented with 4% of cerebrospinal fluid (CSF). The data shown are mean ± SD of normalized relative quantities (NRQ) obtained from transcript levels. At least three independent samples were used, and four technical replicates were performed from each sample. The CAMHB was used as reference. Statistical significance (*p*< 0.05) was determined was determined by *t* test, one asterisks: *p*< 0.05; two asterisks: *p*< 0.01, and three asterisks: *p*< 0.001.

**Figure S2.**
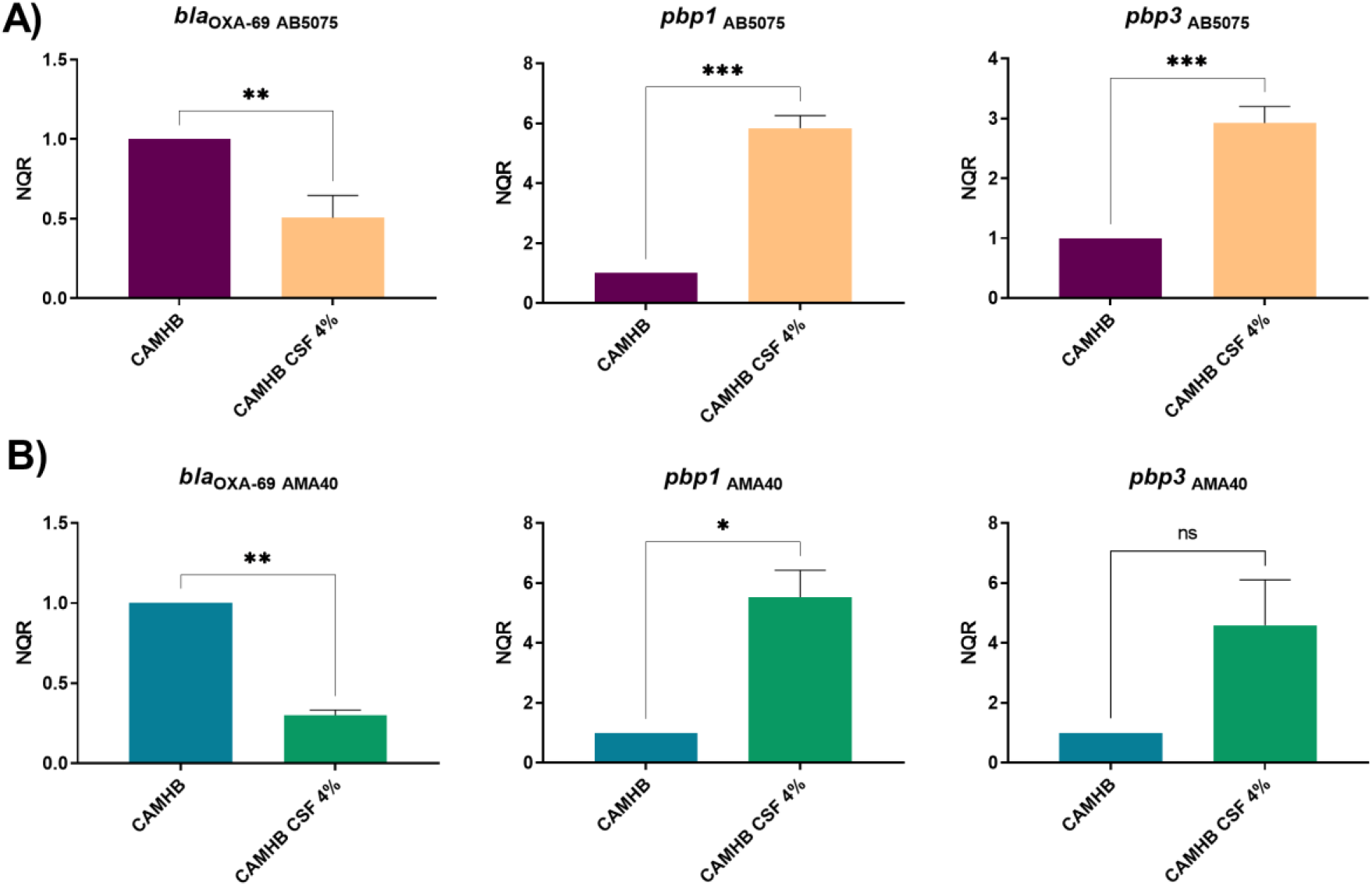
Genetic analysis of β-lactamase and PBP genes of AB5075 (A) and AMA40 (B) *A. baumannii* strains. qRT-PCR of genes associated with β-lactams resistance, expressed in cation adjusted Mueller Hinton (CAMHB) or CAMHB supplemented with 4% of cerebrospinal fluid (CSF). The data shown are mean ± SD of normalized relative quantities (NRQ) obtained from transcript levels. At least three independent samples were used. LB was used as the reference condition. Statistical significance (*p*< 0.05) was determined was determined by *t* test, one asterisks: *p*< 0.05; two asterisks: *p*< 0.01, and three asterisks: *p*< 0.001.

**Figure S3.**
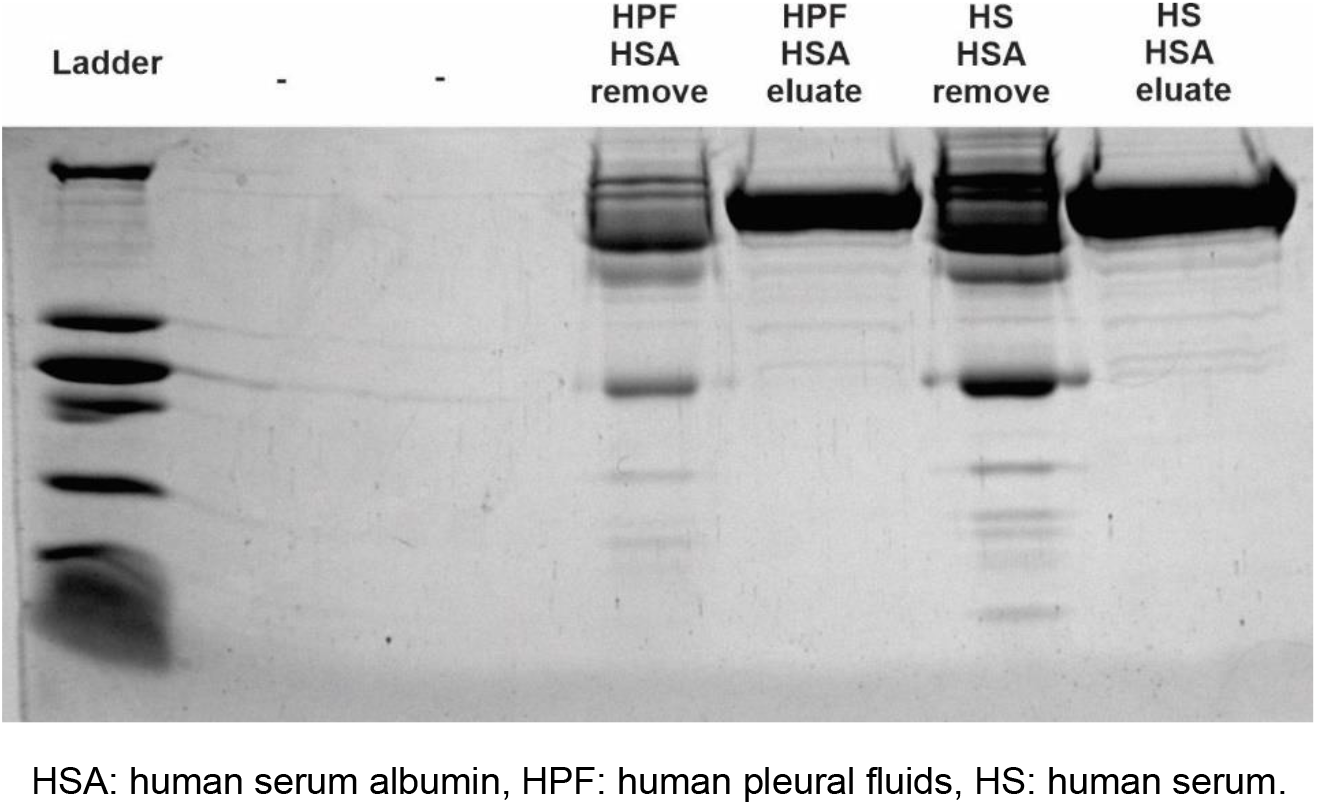
SDS-PAGE of human fluids analyzed using the ProteoExtract® Albumin/IgG Removal Kit (Sigma-Aldrich, MA, United States).

